# Antinociceptive properties of an oral formulation of Δ9-tetrahydrocannabinol in aqueous 2-hydroxypropyl-β-cyclodextrin in female rats

**DOI:** 10.64898/2026.08.04.742765

**Authors:** Farshid Bagheri, Maria Scherma, Elisabetta Murru, Grazia Contena, Sebastiano Banni, Antonio Argiolas, Maria Rosaria Melis, Paola Fadda, Fabrizio Sanna

## Abstract

**Background:** Cannabis derivatives have been reported to possess antinociceptive properties. However, oral delivery is limited by poor bioavailability, stability, and reliability of effects. Previously, we reported an analgesic effect of the aqueous complex Δ9-tetrahydrocannabinol/2-hydroxypropyl-β-cyclodextrin (THC/HPβCD) after intracerebroventricular administration in male rats.

**Methods:** Here, we investigated the analgesic effects of the THC/HPβCD complex after oral administration (0.3 and 3 mg/kg) by the tail flick test after both acute and chronic administration (15 days) in female rats. Locomotor activity and anxiety-like behavior were also evaluated at the same experimental conditions. Moreover, dopamine and glutamate content in the periaqueductal gray (PAG), a key area for the antinociceptive action of THC, were also measured by HPLC.

**Results:** After acute administration, the antinociceptive effect of the complex was seen at 3 but not 0.3 mg/kg THC, with a maximum effect observed at 30 min (MPE 60%). Similar results were obtained after 15 days of treatment, although partially reduced (max MPE 20%). Reductions in locomotor activity with the dose of 3 mg/kg and a slight biphasic effect of the two doses on anxiety-like behavior were also observed. Finally, neurochemical analyses revealed that the dose of 3 mg/kg significantly increased dopamine and glutamate content in the PAG, an effect no longer present after 15 days of treatment.

**Conclusions:** Our results highlight the antinociceptive efficacy of the THC/HPβCD complex also after oral administration, notably higher than that previously seen with other carriers, although with some degree of tolerance after chronic administration. From a translational point of view, these results are relevant for the development of THC-based oral formulations with analgesic properties for the treatment of pain in humans.

**Graphical abstract:** 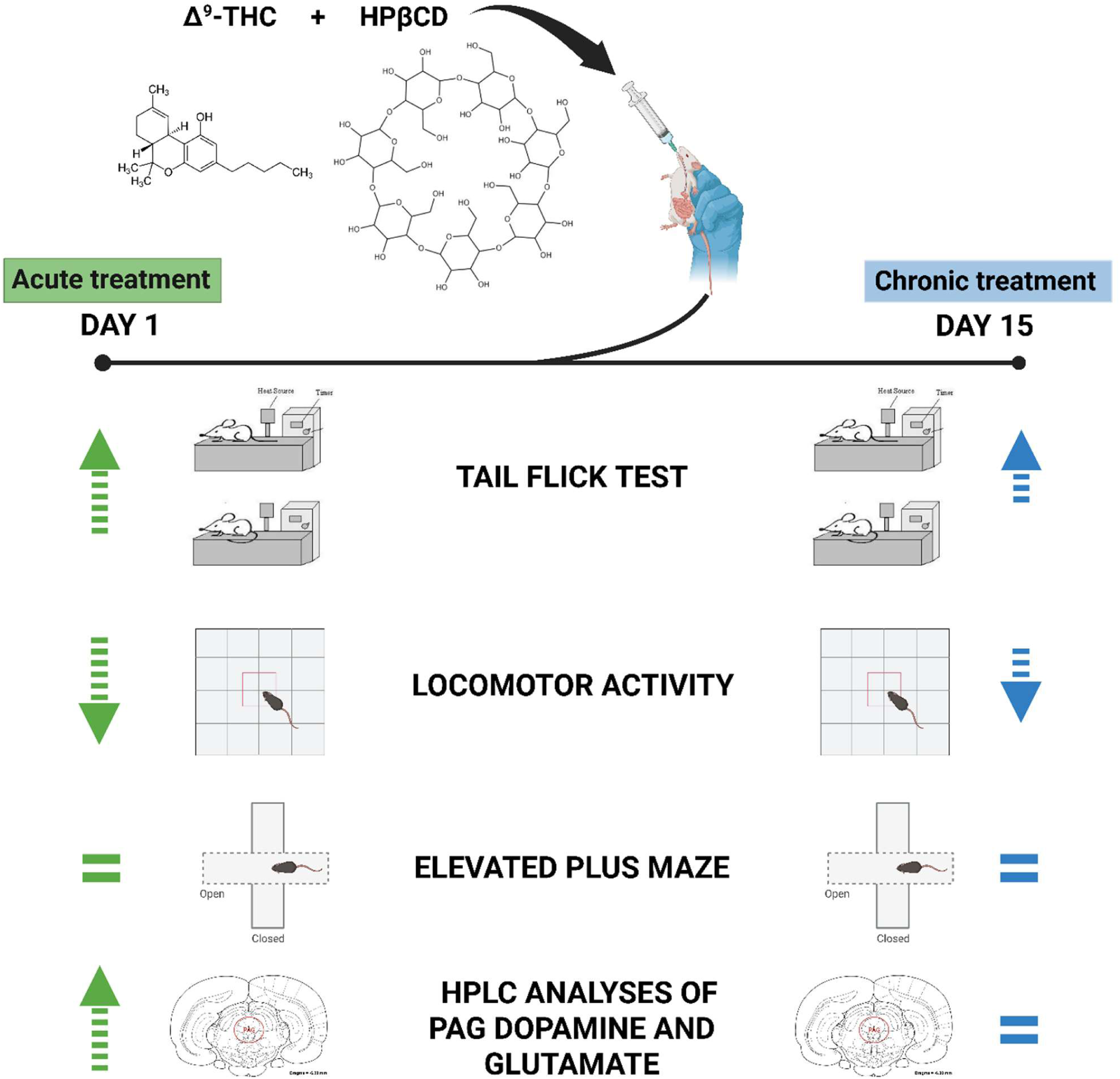

## 1. Introduction

Cannabis derivatives are receiving increasing attention due to their potential therapeutic applications for the symptomatic treatment of pain in different pathological conditions, such as cancer [1], HIV/AIDS [2], neurodegenerative disorders [3], and neuropathic chronic pain [4]. However, the therapeutic use of cannabis derivatives in the clinical practice is limited by several factors. First, these substances are highly lipophilic, and their administration often requires the use of solvents that limit their use in humans [5–9]. Second, oral administration raises issues related to bioavailability, stability, and reliability of effects. When taken orally, Δ9-tetrahydrocannabinol (THC) undergoes extensive first-pass metabolism, and the amount of THC that reaches systemic circulation (bioavailability) is very low (6–20% of the administered dose) [10]. Absorption is also slow, irregular, and variable, even when THC is dissolved in oil. Initial plasma concentrations and peak levels are low and delayed. Moreover, the metabolism of orally administered THC induces the formation of psychoactive metabolites (11-hydroxytetrahydrocannabinol [11-OH-THC] and 11-nor-9-carboxy-tetrahydrocannabinol [THC-COOH]), whose presence in plasma is detectable for several hours. This pharmacokinetic profile results in delayed and variable absorption and is also limited by side effects such as allergic reactions and psychotropic effects.

Although several aspects of THC’s pharmacokinetics and metabolism, as well as some important pharmacological effects, have been well characterized, studies that directly evaluated the potential for improving its current oral administration in terms of bioavailability, stability, and therapeutic index remain relatively limited. Recent studies have explored the possibility of encapsulating hydrophobic drugs, including cannabinoids, in cyclodextrin-based carriers to improve their solubility and release profiles. Accordingly, cyclodextrins are excellent solubilizers for highly lipophilic substances and are present in formulations already available on the market [11,12]. Cyclodextrins are cyclic oligosaccharides used to increase the aqueous solubility, physical stability, and bioavailability of hydrophobic drugs [13, 14] and have been included by the FDA in the list of inactive pharmaceutical substances considered safe for human use [15]. Among the various cyclodextrins, 2-hydroxypropyl-β-cyclodextrin (HPβCD) is administered to humans via the cerebrospinal fluid in the treatment of Niemann-Pick disease [16, 17]. Recently, it has been used experimentally in the subcutaneous administration of progesterone in humans, in the subcutaneous, intramuscular and intravenous administration of diclofenac [18] or in the oral administration of insulin [19, 20]. However, a critical gap persists specifically regarding THC/HPβCD complexation for oral administration both in humans and in rodent models.

A previous study [21] from our group showed that male rats that received intracerebroventricular (ICV) administration of THC complexed with 2-hydroxypropyl-β-cyclodextrin (HPβCD), exhibited increased tail flick latency (by about 30%), indicating antinociceptive properties by the complex. In particular, rats that received the dose of 135 μg, but not those who received 30 μg, displayed a robust and sustained analgesic effect in the tail flick test, and reduced locomotor activity, compared with rats in the control group. The antinociceptive effect induced by ICV administration of THC/HPβCD at the dose of 135 μg was of a similar magnitude to that described in other preclinical studies in which comparable doses of THC were administered centrally using DMSO or Cremophor as solvent [22–24]. To note, our results also confirmed the safety profile HPβCD as a carrier for the delivery of THC in rats; in fact, no adverse or toxic effects were detected both during and after treatment.

As regards the putative brain area(s) mediating the analgesic effects of THC, a broad literature suggests, among others, a key role for the periaqueductal gray (PAG) [23, 25], where the endocannabinoid system interacts with other neurotransmitters such as opioids, glutamate and GABA to modulate pain perception [26–28]. In particular, glutamate and GABA have been reported for exerting bidirectional effects in this brain area on nociception, with glutamate inhibiting it and GABA acting in the opposite direction [29]. The PAG also contains dopaminergic neurons [30] involved in the modulation of pain perception [31], supporting the notion that PAG dopamine mediates antinociception. Accordingly, the dopamine agonist apomorphine when injected into the PAG induces antinociceptive effects [32], while 6-OH-DA lesions of these DA neurons as well as pharmacological blockade of dopamine receptors prevent and/or abolish these effects, although some inconsistency about the subtype of dopamine receptors involved has been reported [33, 34].

Based on these previous findings, the aim of the present study is to investigate the analgesic properties of the THC/HPβCD complex after oral administration in female rats with the tail flick test. Female rats were used instead of male rats because preclinical studies demonstrate that several cannabinoid receptor agonists that act at the CB1 receptor, including THC, are more potent in female rats compared to male rats on various nociceptive tests (i.e. thermal or mechanical) [35], and that tolerance to THC develops more rapidly and to a greater extent in female rats compared to male rats [36]. These considerations further support the relevance of investigating THC effects in females, also in light of emerging evidence indicating that cannabinoid exposure during sensitive physiological conditions, such as pregnancy, can lead to long-term and sex-dependent metabolic and neurobiological alterations in the offspring [37].

To this aim, female rats were orally treated under an acute (only once) or chronic (15 days) protocol with 0.3 or 3 mg/kg THC dissolved in a 22% aqueous solution of HPβCD, or 22% HPβCD alone [21] and tested in the tail flick test in order to assess pain sensitivity at subsequent time points (i.e., 30, 60 and 120 min). Locomotor activity and anxiety-like behavior were also evaluated after both acute and chronic treatment. Finally, in order to investigate the potential involvement of the periaqueductal gray (PAG) in the antinociceptive effects observed, the PAG tissues from the same rats used for the tail flick experiments were processed and analysed by means of high-pressure liquid chromatography (HPLC) for the determination of dopamine and glutamate content.

## 2. Materials and Methods

### 2.1 Chemicals and Reagents

Synthetic Δ9-tetrahydrocannabinol (THC, dronabinol) was acquired from THC Pharm GmbH (Frankfurt, Germany). Hydroxypropyl-β-cyclodextrin (HPβCD) was obtained from Sigma–Aldrich (St. Louis, MO). All other reagents, unless otherwise specified, were of the highest available purity.

### 2.2 Preparation of formulations

A concentrated stock solution of THC (200 mg/mL) was prepared by dissolving 1 g of THC in 5 mL of ethanol at approximately pH 6.8. Subsequent complexation of THC with HPβCD was performed using a previously described methodology [21]. By subsequent dilution steps, three final THC-HPβCD complex solutions were prepared at nominal doses of 0, 0.3, and 3 mg/kg. The vehicle solution was an aqueous 22% (w/v) HPβCD solution with 0.5% ethanol (v/v), as per the ethanol concentration in the 3 mg/kg THC/HPβCD solution. When made, all solutions were kept equilibrated at 37°C with constant 100 rpm shaking for 24 hours prior to use. Doses used were chosen based on previous literature [see for example, 38, 39, 40, 41, 42]. The formulations of THC in aqueous HPβCD were analysed using Fourier transform infrared (FT-IR) spectroscopy to study intermolecular interactions and molecular integrity. The comprehensive method has been already reported [21].

### 2.3 Animals

Young adult female Sprague Dawley rats bred in the animal facility of the University of Cagliari (CeSASt), weighing between 225–275 gr., were used in all the experiments. Eighty-four rats were used in total. Sixty rats were used for the tail flick and locomotor activity experiments. Two of them were excluded before the beginning of the experiments due to gross motor alterations. Twenty-nine rats were allocated for acute treatment and the other 29 for chronic treatment experiments (in both cases: vehicle n = 9; 0.3 mg/kg THC n = 10; 3 mg/kg THC n = 10). The elevated plus maze test was performed in a separate group of 24 female rats (n=8 for each treatment group at the same doses as reported above). In this case, the same female rats were used for the behavioral assessment both after the acute (i.e., after the first treatment) and chronic (i.e., after 15 days of treatment) protocol of administration.

For seven days before the experimental procedures rats were acclimated to the environmental conditions of the animal facility. To minimize stress from handling during the experiments, each rat was handled daily and familiarized with the experimental rooms and procedures as specified below. The rats were housed in groups of four per cage with access to food and water ad libitum and maintained at 22 ± 2°C, with 60% humidity and a 12-hour light/dark cycle (lights on from 08:00 to 20:00). Cages were equipped with standard plastic and wooden environmental enrichment. According to the 3R principles governing animal experimentation, all possible efforts were made to minimize animal suffering and reduce the number of animals used. Additional details about the experimental procedures are reported in the dedicated sections. All experimental procedures were performed in strict accordance with the ARRIVE guidelines, with the European Directive 2010/63/EU and Italian legislation (D.L. March 4, 2014, no. 26) (Italian Ministry of Health, Aut. n. 942/2023-PR) and with the policies issued by the Organism for Animal Welfare (OPBA) of the University of Cagliari.

### 2.4 Experimental conditions and drug administration

Before starting the treatments, rats were randomly divided into three cohorts. Two of them were devoted to the tail flick and locomotor activity experiments (acute vs chronic treatment), while a third cohort was used for the anxiety studies with the elevated plus maze test, and each of them was further divided in three treatment groups: a control group (aqueous 22% (w/v) HPβCD solution), and two treatment groups (0.3 mg/kg and 3 mg/kg THC in 22% HPβCD). Randomization was performed with a free “random numbers generator” (https://www.calculator.net/random-number-generator.html). Rats were treated only once (acute condition) or for 15 consecutive days (chronic condition) once a day between 9.00 and 11.00 am and behavioral assessments were performed as specified below. In order to avoid possible intervening confounding factors, rats from the same litter were distributed in all the experimental groups in a counterbalanced manner. Also, the order of treatments and the position of the cages in the facility room was counterbalanced.

For oral gavage, rats were acclimated to handling and the gavage procedure over a period of 7 days prior to the start of the experiments to reduce stress and ensure consistent administration. A flexible gavage propylene needle (18 gauge) (Instech Laboratories, Inc.) was used to administer the solution directly into the gastric cavity. In order to ensure consistency in the oral administration, food was removed from the cages 2 hours before treatment. Each administration session was monitored closely to confirm that the procedure was well-tolerated by rats.

In all the experimental phases (i.e., treatment, experimental procedures, analysis of data), the experimenter was blind to the treatment received by the animals.

### 2.5 Determination of the phase of the estrous cycle

Estrous cycle phase was determined as previously described [43] by morphological inspection of vaginal smears collected by lavage, i.e., by inserting a 200 μl plastic pipette with a smooth silicone tip filled with saline (0.9% NaCl) into the rat vagina to a depth of approximately 2.0 mm. Briefly, the vaginal smear from each female rat was spread on a glass slide, allowed to dry overnight, stained with May-Grunwald-Giemsa stain, and observed under a phase-contrast microscope, using 10x and 40x objectives. The cycle phase was identified by the morphological features of the vaginal smear and the presence of the following 3 cell types: a) round, nucleated epithelial cells, b) irregularly shaped, keratinized cells, and c) small, dark-stained leukocytes. The different ratios between these cells have been used to detect the following phases of the estrous cycle: Dioestrus: predominance of leukocytes; Proestrus: predominance of large, round, nucleated epithelial cells, which may be grouped into layers; Estrous: predominance of large, pink stained, irregularly shaped keratinized cells; Metestrus: leukocytes, epithelial cells, and keratinized cells in equal proportions.

### 2.6 Tail Flick Test

The tail flick latency was measured to assess the antinociceptive effects of the THC-HPβCD formulation [21]. On the day of the experiment rats were transported from their home cages to the experimental room for a 20-minute habituation period before treatment. Thereafter tail flick response was measured at 30, 60, and 120 minutes after treatment. Tail flick response was assessed after one single treatment (acute response) or at the end of the fifteen days of treatment (chronic response). During the procedure, the rat is placed on a specialized apparatus (TSE Systems, Bad Homburg, Germany) and a focused beam of light, heated to 56°C is projected onto its tail, specifically targeting a point approximately 6 cm from the tip. This exposure to radiant heat prompts a tail flick response, which is the rapid withdrawal of the tail from the heat source. The duration between the onset of heat exposure and the tail flick response is measured and recorded as the “tail flick latency”. To minimize the risk of tissue damage, the exposure is capped at a cut-off time of 20 seconds. Moreover, for each rat the tail flick reflex measurement was done paying particular attention in directing the light beam in three different segments of the tail in a counterbalanced order that accounts both for the treatment group and the time point assessed (30, 60, and 120 minutes); this was done to further minimize the risk of tissue damage and/or the emergence of carry over effects (i.e., the potential interference of a previous measurements on the subsequent ones). The latency to tail withdrawal was carefully recorded for each rat, with the final reported value being the average of two consecutive measurements taken two minutes apart. Baseline latency was determined for each rat before treatment to account for individual variability. The percentage of maximum possible effect (%MPE) was calculated using the formula: %MPE = (experimental latency - mean baseline latency)/(cut-off latency - mean baseline latency)*100, and %MPE values were statistically analysed to compare treatment effects over time, as detailed in the Statistics section.

### 2.7 Locomotor Activity

Locomotor activity was measured as previously described [44]. Before the start of the experiments, rats were handled daily for at least one week to avoid handling stress during the experimental sessions. At the end of this period, each rat underwent a 1-h habituation session to prevent the influence of novelty factors related to the experimental procedure and the motility apparatus during the experimental sessions. On the day of the experiment, rats were transported from their cages to the experimental room for a 20-min habituation period and then the treatments were performed. Rats were individually tested for motor activity under standardized environmental conditions (in a soundproof room with a light level of 30 lux) with a Digiscan animal activity analyzer (Omnitech Electronics, Columbus, Ohio). Each cage (42 cm × 42 cm × 63 cm) was equipped with two sets of 16 photocells arranged at right angles to each other, projecting horizontal infrared beams 2.5 cm apart and 2 cm from the bottom of the cage, and an additional set of 16 horizontal beams whose height was adjusted to the size of the animals (20 cm). Horizontal and vertical activity was measured as the total number of sequential interruptions of the infrared beams (counts) in the horizontal or vertical sensors, recorded every 5 minutes, starting immediately after the animals were placed in the cage, for a testing period of 15 minutes, while the centre time indicated the time in seconds spent by the rat in the central part of the cage, recorded at the same time intervals. Locomotor activity was assessed 60 and 120 minutes after the first or fifteenth (i.e., last) drug administration.

### 2.8 Elevated plus maze test

Anxiety-like behavior was assessed by the elevated plus maze test (EPM) [45,46], a test that exploits the natural tendency of rodents to limit their stay in elevated spaces without protection (where they could be easily spotted by a predator). The apparatus is a black polyvinyl chloride maze equipped with two open and two closed arms (12 × 60 cm) that converged on a small central square (12 × 12 cm), which served as starting point, thus reproducing the shape of a plus sign; the apparatus was elevated 50 cm from the floor and placed in the centre of a sound-proof room under 30 lux light conditions. Thirty minutes after the treatment, the rats, tested one at a time, were placed on the central square facing one of the two open arms and left free to explore the maze for 5 minutes. Rats’ behavior was videotaped by a camera positioned above the maze. The number of entries and the time spent in the open sections provide a measure of behaviors related to the animal’s state of anxiety; in particular, the percent of time spent in open arms [time in open arms/(time in open plus in closed arms)*100] and the percent of entries in open arms [number of entries in open arms/(entries in open plus in closed arms)*100] were calculated. The total number of arm entries (open + closed arms) that provides an index of the level of motor activity of the animal was also calculated. The animal was considered inside a specific arm when it had all four paws inside that arm. The EPM was thoroughly cleaned with H_2_O_2_ before each test to avoid olfactory cues.

### 2.9 Tissue collection and processing

Twenty-four hours after the last drug administration rats were sacrificed by decapitation, the brains were quickly extracted, rinsed and positioned in a rat brain matrix. Coronal brain slices of 2 mm as per Paxinos and Watson (2007) rat brain atlas [47], were made using a stainless-steel blade. Subsequently, the regions containing the periaqueductal gray (PAG), were extracted through the micropunching technique (Supplementary Fig. 1), as previously described [48]. The extracted tissues were weighed and homogenized in 0.1 M perchloric acid (1 mg of wet tissue per 20 µl of 0.1 M HClO_4_) and then centrifuged at 23,000 g for 30 minutes using an Eppendorf 5424R centrifuge (Fisher Scientific, Illkirch, France). Subsequently, the supernatant was filtered through microspin centrifuge tubes equipped with a 0.22-μm nylon filter at 10,000 g for 10 minutes and stored at-80 °C until the day of chromatographic analysis.

### 2.10 HPLC analyses of brain tissues

Dopamine and glutamate were quantified in PAG tissues by HPLC as previously described [49]. Dopamine was measured by injecting a 20 μL aliquot of the supernatant obtained from homogenates by using HPLC coupled to electrochemical detection using a 4011-dual cell (Coulochem II, ESA, Cambridge, MA, USA). Detection was performed in reduction mode at +350 and −180 mV. The HPLC was equipped with a Supelcosil C18 column (7.5 cm ×3.0 mm i.d., 3 μm particle size; Supelco, Supelchem, Milan, Italy), eluted with 0.06 M citrate/acetate pH 4.2, containing methanol 20 % v/v, 0.1 mM EDTA (ethylenediaminetetraacetic acid), 1 μM triethylamine, and 0.03 mM sodium dodecyl sulfate as a mobile phase, at a flow rate of 0.6 mL/min and room temperature. The sensitivity of the assay was 0.125 nM.

Determination of glutamate concentration in the PAG tissue was carried out in 5 μl aliquots of samples diluted 1:20 in HClO_4_ 0.1M in order to prevent column overload and after pre-column derivatization with orto-phtalaldialdehyde and 2-mercaptoethanol, by HPLC. The chromatograph was equipped with a 15 x 0.4 cm Supelco C18 column, 5 μm particle size, and coupled to fluorescence detection (excitation wavelength: 318 nm; emission wavelength: 452 nm; SFM 25 spectrofluorimeter, Kontron, Milan, Italy), using an automatic injector. The mobile phase was phosphate buffer 0.1 M, pH 6.2 containing methanol 30% v/v and the flow rate was 1 ml/min. The column temperature was maintained at 35 °C. The sensitivity of the assay was 100 nM.

For representative chromatograms of dopamine and glutamate HPLC analyses see Supplementary Fig.2.

### 2.11 Statistics

The number of rats needed to perform the experiments presented here was estimated through a priori sample size calculation by using the software G-Power and based on the primary outcome (i.e., tail flick latency) with the following parameters: alpha = 0.05; 1-beta = 0.85; effect size of medium magnitude f = 0.25. The experimental unit was the single animal. Data are presented as mean values ± SEM of absolute values (e.g., in the case of locomotor activity parameters and of the number of total arm entries in the EPM) or percents (e.g., in the case of the tail flick reflex and of the open arm entries and time in the open arms in the EPM).

Differences in the distributions among the treatment groups of the number of females in the different phases of the estrous cycle were analysed by the Chi-square test. Primary and secondary outcomes were analysed by means of one-or two-way ANOVA with the treatment as between subjects’ factor and time (i.e., timepoint of the test or the length of treatment) as within or between subject’s factor depending on the case. Before performing ANOVA, data sets of each of the different experimental variables were inspected for outliers by Grubbs’ test. Homogeneity of variances among the experimental groups was evaluated with the Bartlett’s or Levene’s test depending on the case. Normality of data distribution was assessed by Shapiro-Wilks test. When ANOVAs revealed statistically significant main effects and/or interactions, pairwise comparisons were performed by using the Tukey’s multicomparison test. In all the other cases, Bonferroni’s corrected multiple t tests were performed. Statistical analyses were all carried out with PRISM, Graph Pad 8 Software (San Diego, USA) with the significance level set at P < 0.05.

## 3. Results

### 3.1 Influence of the estrous cycle on behavioral and neurochemical parameters across the treatment groups

Since there is a significant influence of sexual hormones during the estrous cycle not only in the regulation of sexual behavior but also in several other physiological and behavioral aspects, such as pain perception [50], as well as a different pharmacological activity of cannabis derivatives in the different phases of the estrous cycle [51], it was monitored in order to determine at what phase of the cycle were the females at the time of the behavioral tests and sampling for *ex-vivo* analyses. This allowed us to perform an *a posteriori* evaluation of any effects of the phase of the estrous cycle on the variables under examination. The results obtained (see Supplementary Table 1) indicated that the females were randomly distributed in the different phases of the estrous cycle across the treatment groups and timepoints both for behavioral and neurochemical assessments, reducing in this way the possibility that hormonal factors may have biased the treatment effects on specific experimental groups or conditions.

### 3.2 Effect of acute and chronic treatment with the THC/HPβCD complex on tail flick reflex

The analgesic effects of the THC/HPβCD complex were assessed by the tail flick test (for details, see Material and Methods section) both after acute and chronic (i.e., 15 days) treatment.

As shown in Fig. 1a, while acute administration of the 0.3 mg/kg THC/HPβCD complex was ineffective, that of 3 mg/kg induced a significant increase in tail flick latency at 30 and 60 min, as indicated by the %MPE values. Accordingly, two-way ANOVA revealed a significant effect of treatment (F(2, 26) = 11.90, p = 0.0002; time, F(2, 52) = 19.45, p < 0.0001; and treatment × time interaction, F(4, 52) = 11.21, p < 0.0001). Moreover, Tukey’s post hoc comparisons revealed significant differences between 3 mg/kg THC/HPβCD and vehicle-treated rats both at 30 and 60 min after the treatment (p < 0.0001 and p = 0.0047, respectively), although a time-dependent reduction of the effect was observed over time, with a tendency of the analgesic effect to decrease and disappear at 120 min.

**Figure 1.**
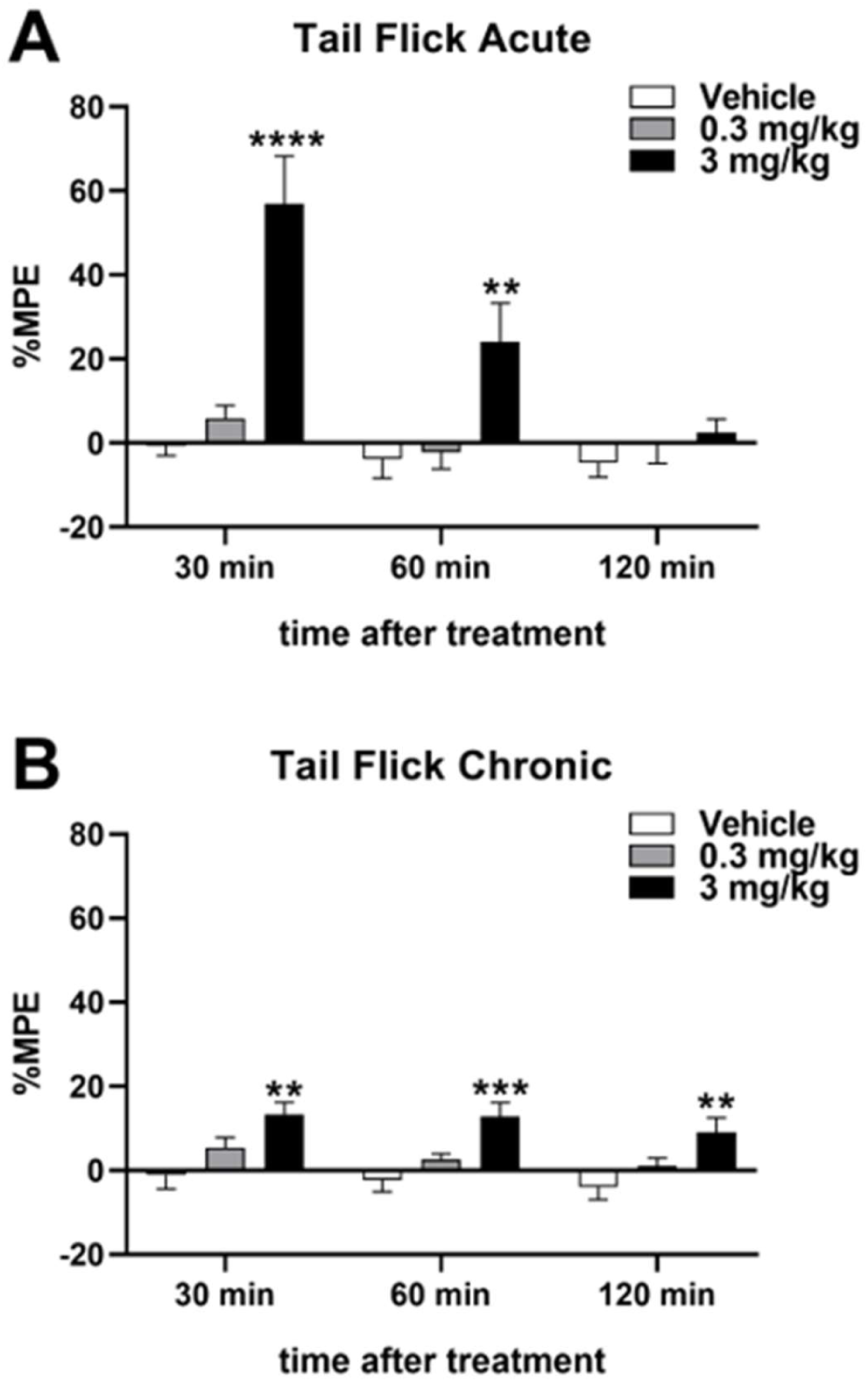
Antinociceptive effects after acute (A) or chronic (B) treatment with the THC/HPβCD complex. Rats were administered either vehicle (22% w/v HPβCD) or formulations containing 0.3 mg/kg or 3 mg/kg of THC complexed with HPβCD via oral administration. The tail flick response was measured at 30, 60, and 120 minutes post-treatment. The results, expressed as %MPE means ± SEM of 9–10 rats/group, were statistically analysed using a two-way ANOVA followed by Tukey’s or Bonferroni’s post hoc comparisons. **: p < 0.01; ***: p < 0.001; ****: p < 0.0001 compared to the vehicle-treated rats. %MPE = (percentage of maximum possible effect, calculated as ([experimental latency – mean baseline latency]/[cut-off latency – mean baseline latency]) × 100).

**Figure 2.**
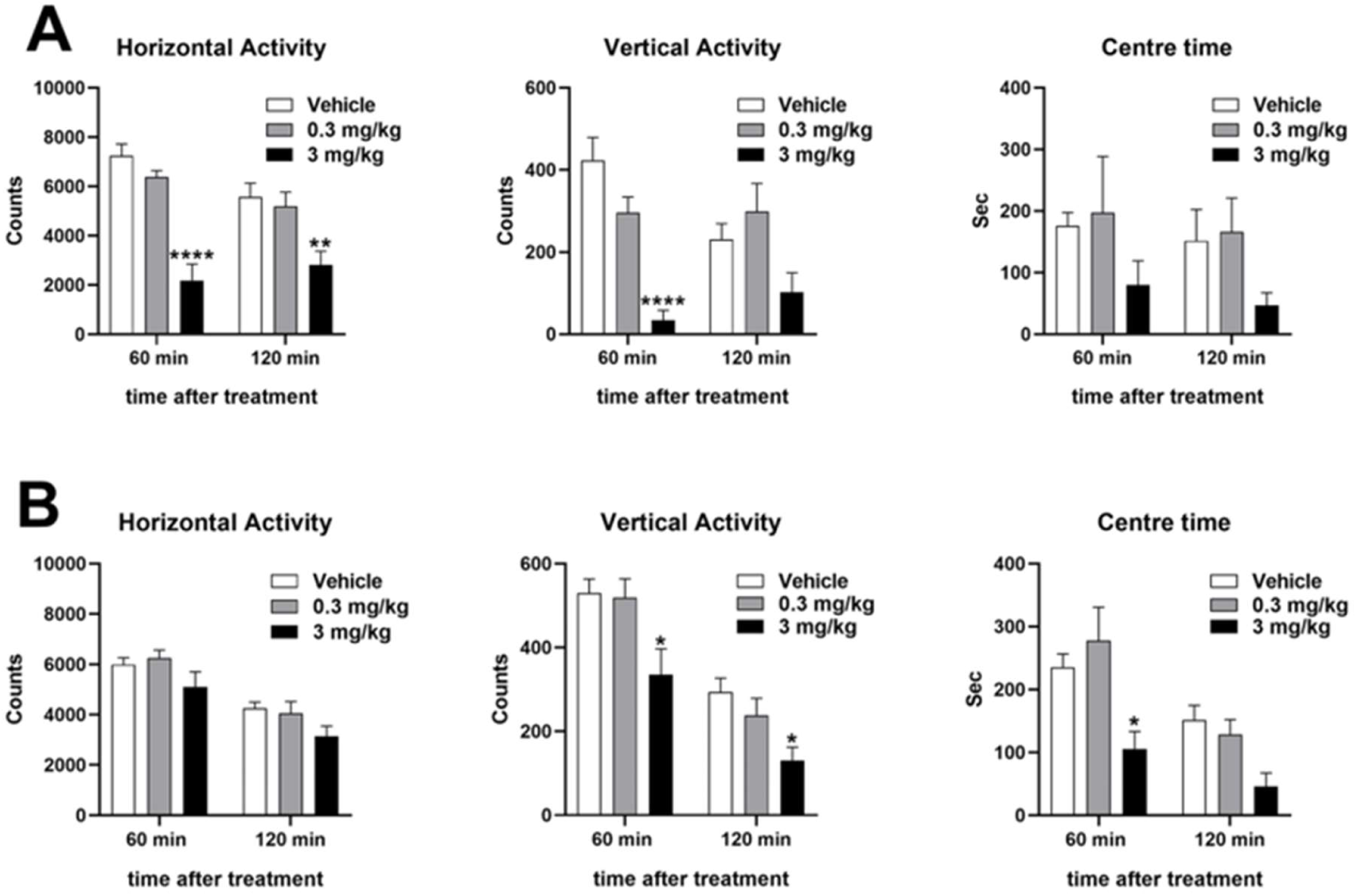
Effect of acute (A) and chronic (B) treatment with the THC/HPβCD complex on locomotor activity. Rats were administered either vehicle (22% w/v HPβCD) or formulations containing 0.3 mg/kg or 3 mg/kg of THC complexed with HPβCD via oral administration. Horizontal, vertical and centre time measurements were taken at 60-and 120-minutes post-treatment. Data are presented as means ± SEM of 6-10 rats/group. Two-way ANOVA followed by Tukey’s or Bonferroni’s post hoc comparisons. *: p < 0.05; **: p < 0.01; ****: p < 0.0001 compared to vehicle-treated rats.

Furthermore, as shown in Fig. 1b, a significant decrease in pain perception was also detected after a 15-day treatment period with the dose of 3 mg/kg. Accordingly, two-way ANOVA revealed a significant effect of treatment (F(2, 26) = 9.25, p = 0.0009 and time, F(2, 52) = 3.41, p = 0.0405), and Bonferroni’s post hoc comparisons revealed significant differences between rats treated with the dose of 3 mg/kg when compared to vehicle-treated rats at 30, 60 and 120 min (p = 0.0014 for 30 min, p = 0.0008 for 60 min and p = 0.0044 for 120 min timepoint, respectively).

However, some differences can be observed between acute and chronic treatment; in particular, lower %MPE values after chronic treatment compared to the acute one at 30 min (13.4% vs 56.8%) but also a longer duration of the effect in the chronic treatment condition, as indicated by the persistence of a significant difference between the dose of 3 mg/kg and the vehicle at 120 min after treatment in this condition.

### 3.3 Effect of acute and chronic treatment with the THC/HPβCD complex on locomotor activity

As shown in Fig. 2a, acute treatment with the THC/HPβCD complex at the dose of 3 mg/kg, but not at the dose of 0.3 mg/kg, resulted in a significant decrease in both horizontal and vertical locomotor activities at 60 and 120 minutes post-treatment. Accordingly, in horizontal activity two-way ANOVA revealed significant effects of time [F(1,15) = 4.561, p = 0.0496] and treatment [F(2,15) = 23.20, p < 0.0001], with a significant time × treatment interaction [F(2,15) = 4.008, p = 0.0403] and Tukey’s multicomparison post hoc test displayed significant reductions of locomotor activity at 60 and 120 min after treatment (p < 0.0001 and p = 0.0026, respectively). Similar results were also obtained for vertical activity, with a significant effect of treatment [F(2, 15) = 11.66, p = 0.0009] and time × treatment interaction [F(2, 15) = 8.814, p = 0.0029] in the two-way ANOVA, and post hoc Tukey’s multicomparisons also indicated significant reductions at 60 min after treatment (p < 0.0001). Finally, regarding the time spent in the centre of the arena, no significant differences were observed between treatment groups, although a pronounced trend to decrease was observed in rats treated with the THC dose of 3 mg/kg at both 60 and 120 min after treatment.

As depicted in Fig. 2b, similarly to what was seen after acute treatment, the 15-day chronic treatment led to a significant reduction in locomotor activity, in particular the vertical activity, and in the time spent in the centre of the arena at the dose of 3 mg/kg, but not at the dose of 0.3 mg/kg. Accordingly, two-way ANOVA detected the following significances: horizontal activity [time: F (1, 26) = 112.8, p < 0.0001]; vertical activity [time: F(1, 26) = 101.0, p < 0.0001; treatment F(2, 26) = 6.41, p = 0.0054]; centre time [time: F(1, 26) = 36.55, p < 0.0001; treatment F (2, 26) = 6.52, p = 0.0051]. Moreover, Bonferroni’s post hoc comparisons detected a significant difference between vehicle and 3 mg/kg treated rats in vertical activity at both 60 and 120 min (p = 0.0071 and p = 0.0278, respectively) and a significant decrease in the time spent in the centre of the arena in rats treated with 3 mg/kg THC compared to vehicles during the first 60 min after treatment (p = 0.0151).

### 3.4 Effect of acute and chronic treatment with the THC/HPβCD complex on anxiety-like behavior

As reported in Fig. 3a, the administration of the THC/HPβCD complex had only a slight effect on anxiety-like behavior as assessed by the EPM test. Accordingly, two-way ANOVA revealed a significant effect of treatment [F(2, 21) = 3.912, p = 0.036] for the percent of time passed in open arms, but Bonferroni’s analyses did not detect any significant differences between the groups treated with 0.3 and 3 mg/kg and vehicle-treated rats. A similar, but not significant, trend was also detected by two-way ANOVA in the percent of entries in the open arms [F(2, 21) = 2.722, p = 0.0889] (Fig. 3b). Finally, no significant differences based on treatment groups and/or type of treatment were detected by two-way ANOVA in the total number of arm entries, a reliable index of the level of motor activity (Fig. 3c).

**Figure 3.**
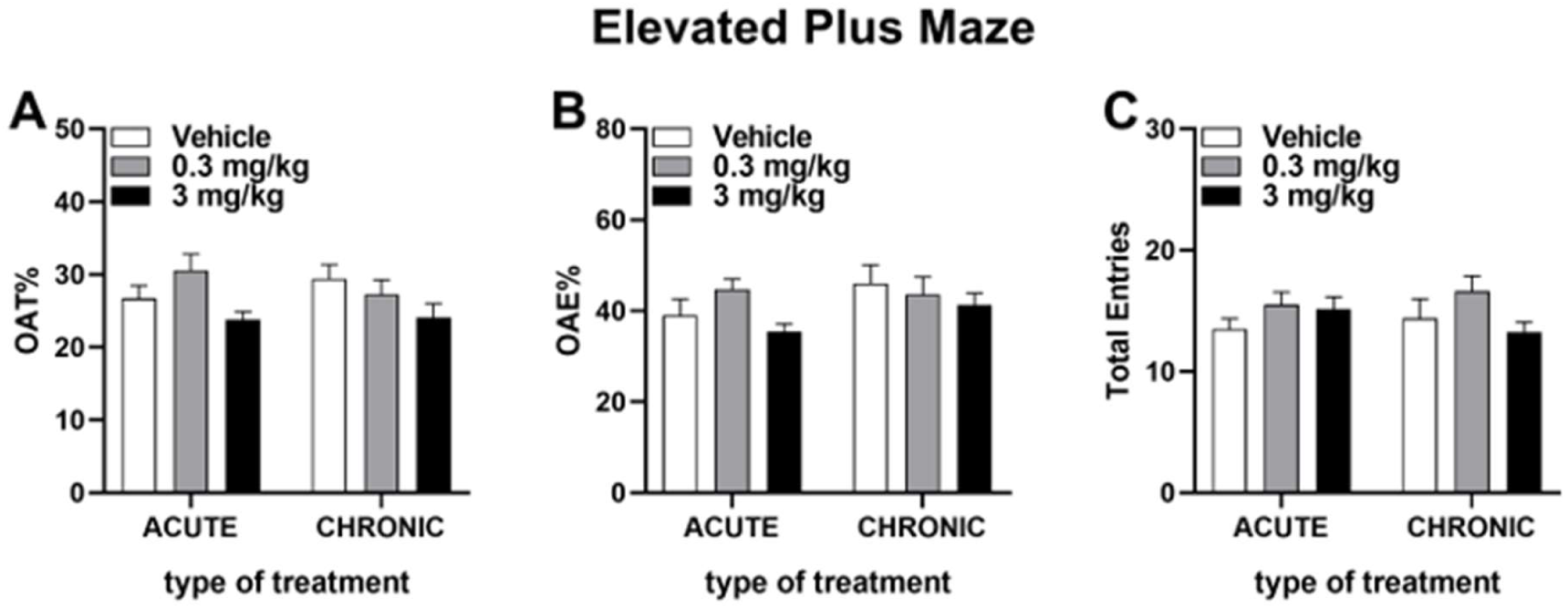
Percent of time spent (A) and percent of entries (B) in open arms, and total number of arm entries (C) in the Elevated plus maze test following acute and chronic treatment with the THC/ HPβCD complex. Rats were administered either vehicle (22% w/v HPβCD) or formulations containing 0.3 mg/kg or 3 mg/kg of THC complexed with HPβCD via oral administration. Percent of time (OAT%) and entries (OAE%) in open arms, and total number of arm entries (open + closed) were taken 30 minutes post-treatment. Data are presented as means ± SEM of 8 rats/group. Two-way ANOVA followed by Bonferroni’s post hoc comparisons.

### 3.5 Effect of acute and chronic treatment with the THC/HPâCD complex on the dopamine and glutamate content of the periaqueductal gray

As reported in Fig. 4, the periaqueductal gray (PAG), a critical brain region for pain perception, showed significant differences among experimental groups in dopamine (DA) and glutamate (GLUT) tissue concentrations following acute but not chronic THC/HPβCD treatment (one-way ANOVA [F(2, 15) = 4.36, p = 0.0322] and [F(2, 15) = 4.061, p = 0.039], for DA and GLUT, respectively). Specifically, acute administration of the THC/HPβCD complex at the dose of 3 mg/kg, but not at the dose of 0.3 mg/kg, significantly elevated DA and GLUT concentrations in the PAG tissue 24 hr after treatment compared to the control condition (+54%, p = 0.0262 and +33%, p = 0.0431, respectively). The effect was dependent on the administration protocol, as it was no longer detectable in rats treated chronically.

**Figure 4.**
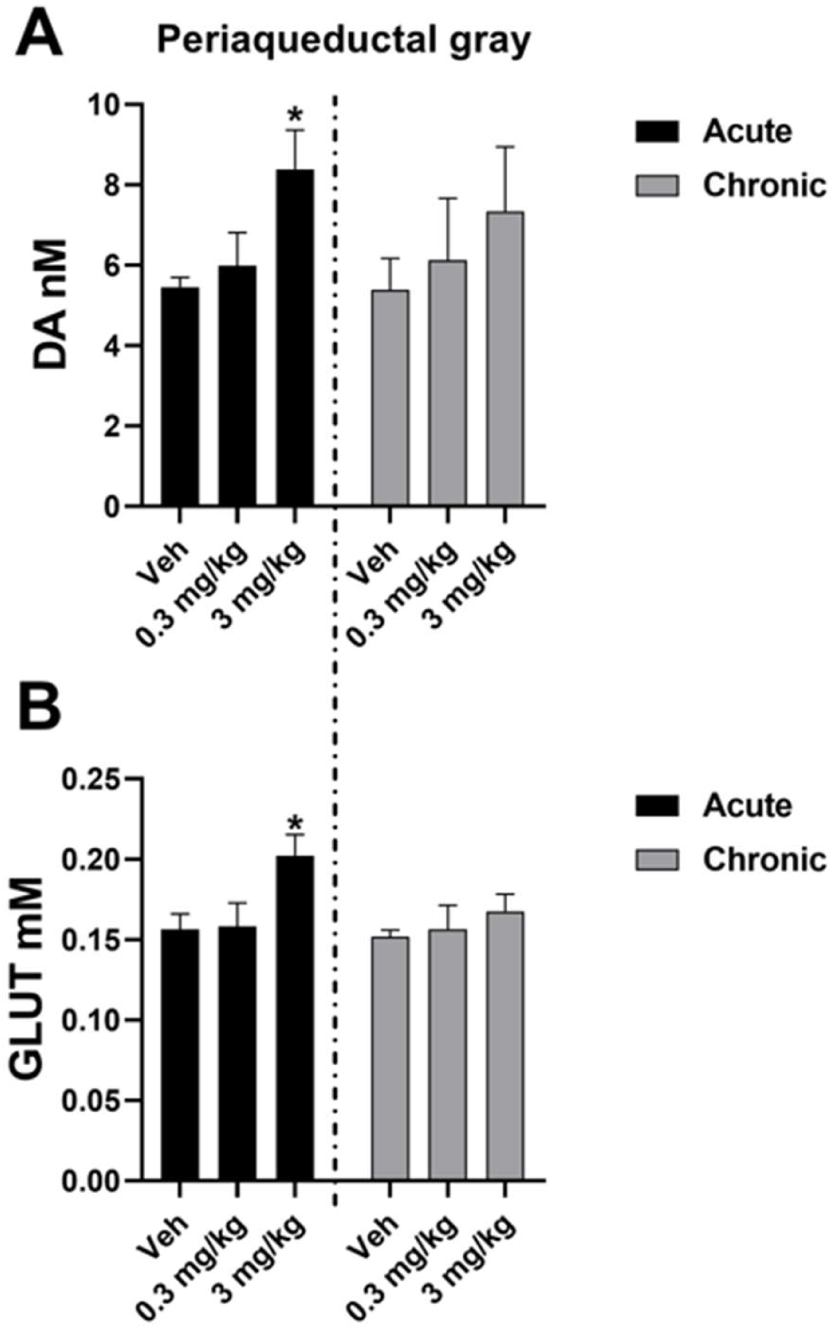
(A) Dopamine (DA) and (B) glutamate (GLUT) concentrations in the periaqueductal gray (PAG) following acute and chronic treatment with the THC/HPβCD complex 24 hours after treatment. Data are means ± SEM of 6-10 rats/group and analysed using one-way ANOVA followed by Tukey’s multicomparisons. *: p < 0.05 compared to the corresponding vehicle-treated group.

## 4. Discussion

This study shows for the first time the antinociceptive effects of a THC/HPβCD complex administered orally in female rats as assessed by the tail flick test, a preclinical model extensively used for the evaluation of the analgesic properties of drugs. The effect was observed at the dose of 3 mg/kg but not at the dose of 0.3 mg/kg, and it was time-dependent with a maximum effect observed 30 min after the administration. Moreover, the analgesic effect was observed both after acute administration (only once) and after 15 days of repeated treatment. However, while the analgesic effect of acute treatment disappeared at 120 min, this was still present at 120 min after 15 days of treatment, although reduced (%MPE change from 60% after acute to 20% after chronic treatment). Other than the analgesic effect, we also observed a significant reduction in both the horizontal and vertical locomotor activity, observed also in this case at the higher dose tested but not at the lower and both after acute and chronic treatment; however, similarly to what was seen in the tail flick test, after chronic treatment the effect was of lower magnitude when compared to that observed after acute administration. Finally, a slight but significant effect of the THC/HPβCD complex was also observed on the anxiety-like behavior. Beyond these behavioral effects, we also observed significant differences in dopamine and glutamate content in the PAG tissues from the rats acutely treated with the THC/HPβCD complex compared with those treated with the vehicle (HPβCD 22% w/v).

Considering the inherent pharmacokinetic and pharmacodynamic complexities of THC, as discussed in the Introduction, the oral delivery pathway poses notable challenges, including substantial first-pass metabolism, unpredictable absorption, and variable onset of action, which have thus far been addressed only superficially by strategies based on the preparation of THC inclusion complexes in substances such as cyclodextrins (HPβCD is one of the most used for this purpose), and the oral administration of these complexes. The ability of cyclodextrins to form inclusion complexes is due to the ability of these oligosaccharides to assume the shape of a truncated cone with a hydrophilic exterior surface and a lipophilic interior cavity, that promotes encapsulation of poorly water-soluble molecules. The size of the interior cavity is suitable to permit formation of 1:1 inclusion complexes with many lipophilic drugs, allowing them to be carried easily across tissues, including the blood brain barrier, and reach target sites to exert their action [52]. Although some studies have shown that HPβCD can significantly improve the aqueous solubility of THC, thus facilitating potential improvements in oral absorption, these results remain largely limited to preliminary proof-of-concept studies, often performed in vitro or using routes of administration that do not reflect typical less invasive therapeutic approaches. For example, formulations using HPβCD for sublingual or parenteral administration in rodents have provided useful insights into the potential of cyclodextrin complexes to improve THC pharmacokinetics. However, these alternative methods do not reproduce the challenges that arise when cannabinoid-derived compounds pass through the gastrointestinal tract, undergo extensive hepatic metabolism, or encounter various enzymatic and transport pathways along the gastrointestinal tract. Consequently, their translational value for oral therapy remains limited. Furthermore, recent literature, although rich in information on cyclodextrin-mediated solubilization, tends to focus on other pharmaceutical active ingredients rather than THC, whose lipophilicity and sensitivity to oxidative degradation require specific complexation strategies.

In a previous work from our group, we reported the ability of the THC/HPβCD complex in inducing analgesia (i.e., increased tail flick latencies) after direct injection in the lateral ventricle of male rats [21]. The present results support the notion that HPβCD is a suitable carrier for the systemic delivery of THC, also through the oral route. This finding can be of high relevance to the development of easy and safe delivery strategies. More important, the magnitude of the effects reported here after oral administration is very similar to that observed after THC administration through other routes such as intraperitoneal and/or subcutaneous or the use of other carriers such as DMSO, ethanol or Cremophor, all of them unsuitable for human systemic administration.

For instance, Lichtman and Martin (1991) [22] observed an analgesic effect in the tail flick apparatus in male rats, after both intravenous and intrathecal administration of THC (3 or 10 mg/kg) dissolved in ethanol-emulphor-saline. Similarly, Tseng and Craft (2001) [53] observed significant effects in the tail withdrawal test in male and female rats with intraperitoneal administration of 10 mg/kg THC in ethanol-emulphor-saline, mainly within the first hour after the treatment. Lazzari et al. (2010) [38] reported significant antinociceptive effects in the tail flick test (25-30% MPE within the first 60 min after the treatment) when 10 mg/kg THC were dissolved in the microemulsion Solutol HS15 and intraperitoneally injected but not in the ethanol-Cremophor-saline solution. However, employing the same ethanol-emulphor-saline carrier, the subcutaneous route required a massive dose of THC (150 mg/kg) to obtain %MPE values of about 60% in male mice 30 min after the administration [54]. In another study [55], the intravenous administration of THC in a mixture of Cremophor/ethanol/water to male mice led to significant increases of 40-60 %MPE in the tail flick latency at doses of 1-2-4-8 mg/kg, 20 min after the administration. However, the antinociceptive properties of Cremophor alone have been shown [56], which questions the suitability of this carrier in analgesia studies. Another study [57] investigated the antinociceptive properties of oral THC dissolved in ethanol observing an increased tail flick latency (by about 60% MPE) 30 min after the administration of a dose of 100 mg/kg. In a more recent study [39] oral administration of THC in sesame oil to male and female rats increased tail flick latency of about 50% MPE in both females and male rats at doses of 5.6 mg/kg or higher (up to 20 mg/kg). However, at variance from our results, this effect was observed not before than 120 min after the treatment, and lasted for at least 300 min, pointing out differences related to the specific carrier employed not only in the efficacy of the drug but also in the temporal window in which the effect is observed. Moreover, the increased tail flick latency at 3 mg/kg was much lower than that reported in the present study (30 vs 60% MPE) and did not reach significance, indicating the higher efficacy of the THC/HPβCD complex in increasing the tail flick latency.

Overall, and regardless some specific differences in experimental methodology, the magnitude of the antinociceptive effects observed with the THC/HPβCD complex is similar, and often superior, to that observed in previous studies employing other carriers and/or more invasive routes of administration. HPβCD’s ability to enhance solubility and potentially improve the bioavailability of lipophilic molecules like THC is evident here, offering a notable improvement over other carriers in terms of efficiency, efficacy and rapidity of drug delivery effects. It should be noted that in the present study we did not measure the ability of HPβCD per sé to modify the response in the tail flick apparatus, for instance, by comparing the tail flick reflex of HPβCD-treated rats with that of rats treated with other vehicles (e.g., saline, distilled water, DMSO and so on). However, this possibility, although cannot be ruled out, is unlikely also considering previous reports showing no effect by cyclodextrins per sé in several models of analgesia [58, 59].

Other than the antinociceptive effects, we also observed that both acute and chronic administration of the THC/HPβCD complex resulted in significant reductions in horizontal and vertical locomotor activity. This effect on locomotor activity was observed at the dose of 3 mg/kg but not at the dose of 0.3 mg/kg, and consistent across treatment durations, although a partial reduction in its magnitude was observed after chronic treatment compared with the acute one. The inhibitory effects of THC on locomotor activity have been broadly and consistently reported in literature [60]. For instance, Tseng and Craft (2001) [53] observed a decrease in motor activity with intraperitoneal administration of 3 and 10 mg/kg of THC dissolved in emulphor/ethanol/saline, mainly within the first 120 min after the treatment, in both male and female rats, although with important sex differences. Similarly, oral administration of THC in sunflower oil at the dose of 10 mg/kg decreased locomotor activity in male rats 120 min after administration, whereas the same dose was unable to modify motor activity after subcutaneous injection [40]. In another study [39], oral delivery of THC at the doses of 5.6-20 mg/kg in sesame oil showed a biphasic effect, with increases of motor activity at 30 min and a decrease at 270 min after the treatment, also in this case with some sex-related differences. However, in our study, the inhibitory effect on locomotor activity was already seen within the first hour after the treatment at the dose of 3 mg/kg and lasted for 120 min, further demonstrating the efficacy of HPβCD in oral delivery of THC, compared with the studies cited above, also in inducing a significant suppression of motor behavior. As noted above, and similarly to what observed for the antinociceptive effects, the magnitude of the suppressive effects of the THC/HPβCD complex on motor behavior was lower after chronic treatment compared to acute administration indicating some degree of tolerance by repeated exposure. The similar trends in the motor and antinociceptive effects of the complex raise the possibility that the increased latencies in tail flick supporting the antinociceptive effects of the complex could be due to the suppressive effects of the complex itself on motor activity. However, this is unlikely, being the tail flick reflex primarily a nociceptive spinal reflex arc. In fact, the essential neuronal circuit is entirely located within the spinal cord, and its execution is independent of the animal’s level of voluntary motor activity or state of consciousness, making it a fundamental model in pain pharmacology [61–63]. Nevertheless, the inhibitory effects of the THC/HPβCD complex on locomotor activity at the same doses of its antinociceptive effects, raises some issues related to the safety of its use for the treatment of pain in humans, revealing that a strict regulation for its use is required in those contexts in which high levels of attention are required such as in schools, workplaces, while driving in the traffic, and so on.

The anxiogenic/anxiolytic effects of THC have been traditionally the object of a long debate and represent a sort of keystone to better understand the therapeutic potential of THC and other cannabis derivatives. The debate is mainly based on the complexity, and somehow partial inconsistency, of the biphasic effects observed, with anxiolytic effects by relatively low doses of THC but anxiogenic effects with higher doses [64, 65]. However, the picture is further complicated by the fact that different carriers and/or routes of administration tend to deeply interfere with the effects observed as well as the species used (e.g., mice vs rats) and sex [66]. Rubino et al. (2007) [67] reported significant anxiolytic effects of intraperitoneal administration of THC at doses between 0.075-1.5 mg/kg dissolved in Cremophor, ethanol, and saline (1:1:18) in male rats 30 minutes before the EPM test, and a similar anxiolytic effect was also observed by Fokos and Panagis (2010) [68] after intraperitoneal administration of 0.5-1.0 mg/kg THC in DMSO, Cremophor, saline. In contrast, Perez-Valenzuela et al. (2024) [69] did not observe any anxiolytic effect in their male and female rats treated with 0.3 mg/kg THC administered orally through edible chocolate 90 min before the test. In another study [70], 2.5 mg/kg THC in emulphor, ethanol, and saline (1:1:18) given intraperitoneally 30 minutes before the test significantly increased anxiety parameters as assessed by the EPM, but in the study of Gom et al. (2025) [71], 5 mg/kg edible THC (mixed in peanut butter) was unable to affect the performance of rats in the EPM when given orally 90 minutes prior of the test. Overall, it seems that the oral administration is less effective in affecting the anxiety status (both in anxiolytic and anxiogenic direction) when compared with the intraperitoneal route of administration, also evidencing the apparent safety of our carrier in vehiculating well-recognized THC-related behavioral effects on anxiety. However, these comparisons are further complicated by substantial differences not only in the routes of administration and carriers used but also in the timing of the experimental protocols. Finally, similarly to what discussed for the antinociceptive effects of the complex, one could raise the possibility that its motoric effects could have influenced the anxiety-like behavior in the EPM test. However, this is unlikely considering that no differences were detected in the total number of arm entries, a reliable index of motor activity, based on treatment groups and/or length of treatment, a finding that lays down against this possibility.

Regardless of the specific pharmaceutical formulation, as pointed out above, after THC administration important sex-related differences are found in pharmacokinetics, and in neurochemical and behavioral aspects, both in preclinical and human studies. These differences are characterized by higher responses in females, in terms of lower doses needed and long-lasting effects, compared to males [72–74]. These sex-related differences, which are secondary to differences in cannabinoid metabolism, cannabinoid receptor expression and influence of ovarian hormones including estradiol and progesterone [51], have been also found to occur for the antinociceptive, motor and anxiety-related effects of THC [50, 66]. The higher sensitivity of females to the effects of THC (and other cannabinoids) and the potential risk of severe long-term negative effects for the offspring if these substances are taken during pregnancy [37] are some of the main reasons that led us to start the present investigation in females. However, this point should be seen as a limit of the present study and experiments involving both sexes are warranted to investigate the potential sex-related differences to the THC/HPβCD complex, both in terms of translational relevance and occurrence of unwanted side effects.

In the present study, significantly higher dopamine and glutamate contents were found in the PAG of rats treated with the dose of 3 mg/kg of the THC/HPβCD complex compared to vehicle treated rats 24 hours after acute administration, while the same effects, although still present, appeared blunted (and not more significant) after 15 days of repeated treatment. The parallelism in the differences in the tail flick response and dopamine and glutamic PAG content between acutely and chronically THC/HPβCD-treated rats raises the possibility that these two neurotransmitters are at least in part mediating the antinociceptive effects of THC at the PAG level (see the Introduction). However, although previous studies reported a direct interaction of the endocannabinoid system with dopamine in regulating pain and analgesia both at central and peripheral level [75–78], no one to date focused on the PAG. Additional experiments are needed to test directly this hypothesis. Yet, in light of the inhibitory effect of the THC/HPβCD complex on motor activity, the involvement of other brain areas in which dopamine-endocannabinoid interactions are present, cannot be ruled out [79, 80]. As regards a putative role of PAG glutamate as a mediator of the antinociceptive effects of the THC/HPβCD complex, it is known that the activation of CB1 receptors in the PAG leads to antinociception by inhibition of local GABA release, and consequent increase of glutamate release [29], which in turn activates the metabotropic glutamate receptors type 5 (mGlu5). The activation of these mGlu5 receptors in the PAG is in turn involved in mediating antinociception in the descending pain pathway through changes on the rostral ventromedial medulla (RVM) ON-and OFF-cells activities [81–83]. Also in this case, further experiments are needed to test this hypothesis.

Weather the increases in dopamine and glutamate concentration found in PAG tissue 24 hr after acute but not chronic 3 mg/kg oral THC/HPβCD administration reflects a supra-spinal antinociceptive action of the THC/HPβCD complex is unknown at present. Although the tail flick response is classically defined as a spinal reflex, several studies suggest that it is also modulated by descending signals from the PAG and other supra-spinal structures [84, 85]. Indeed, electrical stimulation [86] or pharmacological manipulation of the PAG (including dopamine and glutamate neurotransmission modulation) can significantly increase tail flick latency [87, 88], indicating that higher-order pain modulation, not just a spinal cord reflex arc, is being measured with the tail flick test. Thus, it is possible that the tail flick latency increases induced by acute THC/HPβCD reflect a possible correlation with the observed dopamine and glutamate increases occurring in the PAG. Likewise, since such increases do not reach significance after chronic THC/HPβCD, it is also tempting to speculate that this may be correlated to the reduction that occurs in tail flick latencies compared to acute treatment found in this experimental condition, providing a possible neurochemical mechanism for the tolerance that develops to the antinociceptive effect of THC/HPβCD after chronic treatment. Further experiments are required to clarify these points.

Regardless of the mechanisms of action through which the THC/HPβCD complex induces analgesia, our data also show that after 15 days of chronic treatment the magnitude of the behavioral and neurochemical effects is reduced compared to what observed after acute administration. This finding suggests that chronic treatment with the THC/HPβCD complex leads to the development of some degree of tolerance. This is not surprising also in light of the reduction in the recreational and therapeutic effects of THC observed after repeated use/administration both in humans [89, 90] and preclinical models [91, 92], being this reduction mediated mainly by down-regulation of CB1 receptors [93]. Additional experiments are needed to investigate the ability of the THC/HPβCD complex at the doses used here in inducing modifications in the expression of CB1 receptors.

## 5. Conclusions

Overall, our results support the feasibility of oral THC delivery using an HPβCD-based formulation for the induction of antinociceptive and analgesic effects under the present experimental conditions. This finding can be of relevance for the development of safe, effective and non-invasive treatments for pain relief based on cannabinoid phytotherapy. The magnitude of the antinociceptive effects was bigger after acute versus chronic treatment, indicating the development of some degree of tolerance after repeated administration; however, a significant reduction in the response to pain stimuli was still present after 15 days of daily administration. Yet, the analgesic effect was observed at the same dose that induced a significant reduction in locomotor activity, highlighting the need for a strict regulation of its potential use in humans. Moreover, although HPβCDs have proven a safe profile and are currently approved by FDA for human use, studies investigating the safety of the HPβCD/THC complex are warranted, in particular in the case of repeated administration.

Finally, since the study was not designed to directly compare pharmacokinetics or bioavailability across different carriers or administration routes, no definitive conclusions can be drawn in this respect. Given the known influence of oral administration on THC metabolism, including first-pass hepatic conversion to active metabolites and possible lymphatic distribution, further studies specifically addressing these aspects will be necessary to better define the role of the carrier in determining the pharmacological profile of THC.

Nevertheless, oral pharmaceutical preparations based on the THC/HPβCD complex could represent a valid and non-invasive alternative for the treatment of pain in several pathological conditions in humans.

## Author Contributions

Conceptualization, M.R.M., F.S., P.F. and M.S.; methodology, F.S. and M.S.; validation, F.S., M.R.M, P.F., A.A., M.S. and S.B.; formal analysis, F.S., M.S., E.M. and F.B.; investigation, F.B., F.S., M.S., G.C., and E.M.; resources, F.S., M.R.M, A.A., P.F., M.S. and S.B.; data curation, F.B., F.S. and M.S.; writing—original draft preparation, F.B., F.S. and M.S.; writing— review and editing, F.S., F.B., M.R.M, P.F., A.A., M.S. and S.B.; supervision, F.S. and M.S.; project administration, F.S.; funding acquisition, F.S., M.R.M, A.A., P.F., M.S. and S.B. All authors have read and agreed to the published version of the manuscript.

## Statement on animal experimentation

All experimental procedures were performed in strict accordance with the ARRIVE guidelines, the European Directive 2010/63/EU and Italian legislation (D.L. March 4, 2014, no. 26). The animal study protocol was approved by the Ethics Committee of the Italian Ministry of Health (Aut. n. 942/2023-PR, October 31, 2023) and by the Organism for Animal Welfare (OPBA) of the University of Cagliari.

## Data Availability Statement

The raw data supporting the conclusions of this article will be made available by the authors on reasonable request.

## Acknowledgments

We gratefully acknowledge the work of the CeSaSt (Centro Stabulari d’Ateneo) personnel at the University of Cagliari, for animal housing and care. The Graphical Abstract was Created in BioRender. Sanna, F. (2026) https://BioRender.com/b58jtne.

## Abbreviations

The following abbreviations are used in this manuscript: 
THC: Δ9-tetrahydrocannabinol
HPβCD: 2-hydroxypropyl-β-cyclodextrin
PAG: periaqueductal gray
MPE: maximum possible effect

