## Supplementary material for "Antinociceptive properties of an oral formulation of Δ9-tetrahydrocannabinol in aqueous 2-hydroxypropyl-β-cyclodextrin in female rats"

**Research Article**

Division of Neuroscience and Clinical Pharmacology

University of Cagliari,

Cittadella Universitaria di Monserrato,

09042 Monserrato (Cagliari), Italy


**Running title:** Analgesic effect of orally administered THC/HPβCD complex

**Supplementary material**

*Supplementary Table 1.*
Distribution of the females used in the present study in the different phases of the estrous cycle the day of the behavioral experiments and the day of tissues' sampling for *ex vivo* analyses. The exact number of females belonging to the three treatment groups in each of the four estrous phases is reported (Chi square test, all not significant).

| Cohort of female rats | Day of cycle evaluation | Treatment group | P | E | M | D |
|---|---|---|---|---|---|---|
| Acute THC (Tail flick + motor activity) | Day of behavioral tests | 22% HPβCD | 1 | 2 | 1 | 5 |
| | | 0.3 mg/kg THC- HPβCD | 2 | 1 | 2 | 5 |
| | | 3 mg/kg THC- HPβCD | 2 | 2 | 2 | 4 |
| | Day of sacrifice for tissues sampling | 22% HPβCD | 2 | 1 | 1 | 5 |
| | | 0.3 mg/kg THC- HPβCD | 2 | 2 | 1 | 5 |
| | | 3 mg/kg THC- HPβCD | 1 | 2 | 1 | 6 |
| Chronic THC (Tail flick + motor activity) | Day of behavioral tests | 22% HPβCD | 1 | 2 | 1 | 6 |
| | | 0.3 mg/kg THC- HPβCD | 1 | 3 | 1 | 5 |
| | | 3 mg/kg THC- HPβCD | 3 | 3 | 0 | 4 |
| | Day of sacrifice for tissues sampling | 22% HPβCD | 2 | 2 | 1 | 5 |
| | | 0.3 mg/kg THC- HPβCD | 1 | 2 | 2 | 5 |
| | | 3 mg/kg THC- HPβCD | 2 | 3 | 2 | 3 |
| Acute and chronic THC (Elevated plus maze) | Day of behavioral test after acute treatment | 22% HPβCD | 2 | 2 | 1 | 3 |
| | | 0.3 mg/kg THC- HPβCD | 1 | 1 | 2 | 4 |
| | | 3 mg/kg THC- HPβCD | 0 | 3 | 1 | 4 |
| | Day of behavioral test after chronic treatment | 22% HPβCD | 1 | 3 | 0 | 4 |
| | | 0.3 mg/kg THC- HPβCD | 2 | 2 | 1 | 3 |
| | | 3 mg/kg THC- HPβCD | 1 | 3 | 1 | 3 |

Legend: P = proestrus; E = estrous; M = metestrus; D = dioestrus.

*Supplementary Figure 1*

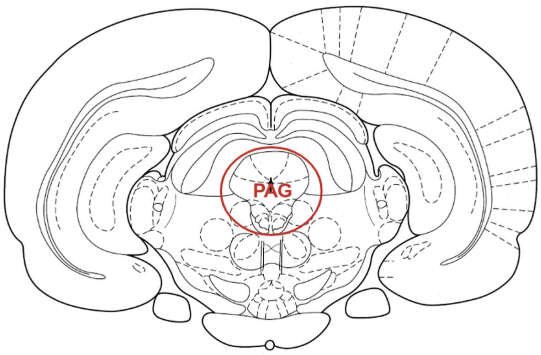


Figure S1. Schematic representation of the brain area interested by the micropunching for the collection of the PAG tissue (evidenced by the red circle) (Bregma = -6.3; Paxinos and Watson, 2007) [47].

*Supplementary Figure 2*

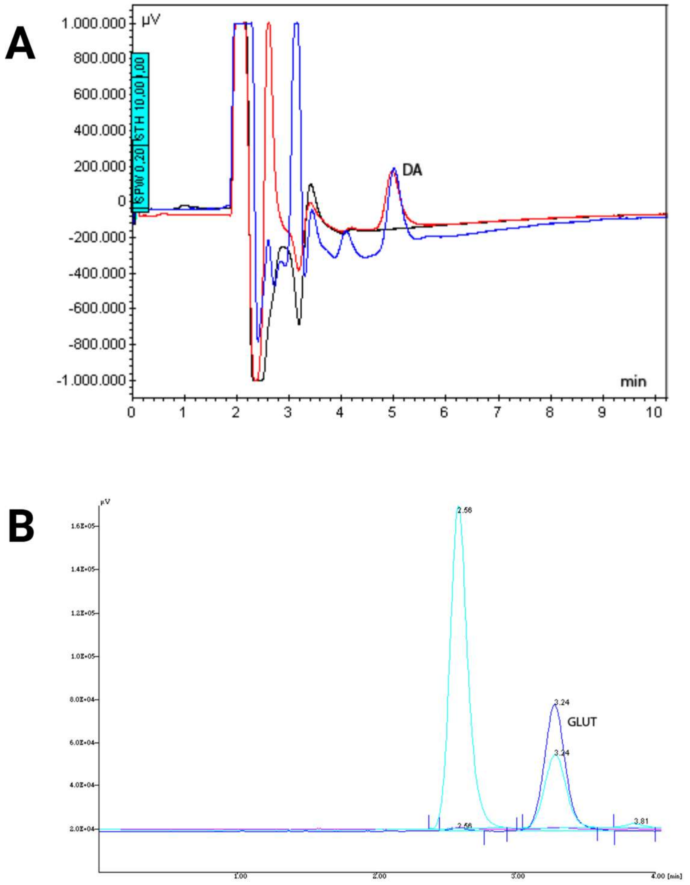


Figure S2. Representative HPLC chromatograms of (A) dopamine (DA) and (B) glutamate (GLUT) concentrations in the PAG tissues from female rats treated with the THC/HPβCD complex or HPβCD 22% (w/v) alone. The black and red lines in the DA chromatogram represent matched blank matrix and a standard sample (5 nM), respectively; the blue line represents a sample from an experimental animal. The purple and dark blue lines in the GLUT chromatogram represent matched blank matrix and a standard sample (10 µM), respectively; the light blue line represents a sample from an experimental animal.
